# Single Molecule Light Field Microscopy

**DOI:** 10.1101/2020.05.20.104802

**Authors:** Ruth R. Sims, Sohaib Abdul Rehman, Martin O. Lenz, Adam Clark, Edward W. Sanders, Aleks Ponjavic, Leila Muresan, Steven F. Lee, Kevin O’Holleran

## Abstract

We introduce single molecule light field microscopy (SMLFM), a novel 3D single molecule localization technique that is capable of up to 20 nm isotropic precision across a 6 *μ*m depth of field. SMLFM can be readily implemented by installing a refractive microlens array into the conjugate back focal plane of any widefield single molecule localization system. We demonstrate that 3D localization can be performed by post-processing 2D localization data generated by common, widely-used, algorithms. In this work we benchmark the performance of SMLFM and finally showcase its capabilities by imaging fluorescently labeled membranes of fixed eukaryotic cells below the diffraction limit.

## 1 Introduction

Single Molecule localization Microscopy (SMLM) has emerged as one of the most popular approaches to super-resolution fluorescence imaging, in part due to the relative simplicity of its experimental implementation (1). In 3D SMLM, axial information is usually obtained at the detriment of both precision and resolution resulting from: a reduction in photon throughput (attributable to additional optical elements), intrinsically higher background from out-of-focus emitters and extended point spread functions (2). We introduce Single Molecule Light Field Microscopy (SMLFM), a simple and highly-efficient 3D super-resolution imaging technique which combines the complementary strengths of SMLM and light field detection to achieve super-resolution imaging throughout a continuous 3D volume.

Numerous approaches for extending the depth of field of single molecule localization microscopy have been developed (3–6). The most successful examples perform single-shot 3D imaging by modifying the shape of the intensity point spread function (PSF) to encode axial information. Astigmatic or rotating Double-Helix Point Spread Functions (DH-PSF) have depths of fields (DOFs) ranging from 0.5 to 4 *μ*m (7, 8). Larger axial ranges have been achieved using other wavefront engineering approaches (9), such as Tetrapod (Saddle-Point) (10) and secondary astigmatism (11). However, in these cases, extracting the super resolved positions of single molecules is generally more challenging than in 2D SMLM as the engineered PSFs cannot accurately be approximated by 2 dimensional Gaussian functions. As a result, more computationally expensive approaches, necessitating phase retrieval (12, 13) or spline fitting (14) to generate finely tuned templates are required. Furthermore, the large spatial footprint of these sculpted PSFs decreases the maximum achievable localization density, significantly decreasing throughput. Another approach, multi-focal plane microscopy (MPM), images a discrete number of axial planes to different lateral positions on one or more detectors (15–18). However, since an emitter located on a particular axial plane is in focus in a subset of these images, whilst contributing background to the others, only a fraction of the total number of detected photons contribute to the precision of each localization.

Light field microscopes built using refractive microlens arrays (MLA) have point spread functions composed of an array of spots, each of which resembles a 2-dimensional Gaussian function. This is also the case for other pupil bisecting methods (19, 20). Each spot remains compact throughout the DOF, which is extended with respect to the widefield case due to the low effective numerical aperture (NA) of each microlens. Hence, existing, optimized, algorithms (21, 22) can be utilized to estimate the location of the centre of each foci with a precision much finer than its width. Information from each 2D localization can be combined to estimate 3D emitter position. The temporal sparsity of SMLM, which limits the probability of overlap between images of distinct emitters, makes it an extremely attractive technique to combine with light field microscopy. We compare two SMLFM configurations tuned to different DOFs by characterising their 3D precision using fluorescent beads in photon regimes encountered in bio-imaging using popular labelling protocols. We also demonstrate efficient detection and localization of single molecules in densely blinking specimens by imaging the membrane of fixed T-cells using both SMLFM configurations, achieving up to 25 3D localizations per frame.

## 2 Light field microscopy

Light field microscopy (LFM) offers single-shot three-dimensional imaging by simultaneously collecting light from a large, continuous depth-of-field. Emitter location is discriminated through wavefront sampling, generally using a refractive microlens array. The MLA partitions a 2D detector into a 2D array of 2D measurements, such that each pixel can be mapped to a 4 dimensional space - known as the light field 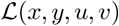. Light field measurements encode both the spatial location and arrival direction of incident photons, where (*x, y*) and (*u, v*) denote spatial and angular co-ordinates respectively. The location of the MLA in the detection path varies but it is generally positioned in either a conjugate image plane or a conjugate back focal (Fourier) plane. The location of the MLA dictates the sampling rates of the spatial and angular co-ordinates of the light field. When the microlens array is placed in an image plane the microlenses themselves sample the spatial domain. Since this configuration results in a loss of spatial resolution, observed most acutely at the image plane (23), the microlens array is optimally located in a plane conjugate to the pupil of the microscope objective for SMLFM (24). This configuration is known as Fourier light field microscopy (25–27).

Each microlens locally apertures the wavefront and generates a focused image, displaced in the direction of, and at a distance proportional to, the average gradient of the apertured wavefront. Hence, not considering aberrations, emitters located on the nominal focal plane are imaged to identical locations in each perspective view. As a result of parallax, axially displaced emitters are imaged to different positions in each perspective view (28). In normalized pupil co-ordinates, the phase in the pupil plane due to an emitter located at (*x*_*i*_, *y*_*i*_, *z*_*i*_) is:

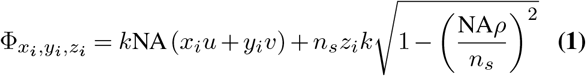

where *ρ*^2^ = *u*^2^ + *v*^2^ = 1 at the pupil edge and 0 at the optical axis, *k* is the free space wavenumber, *n*_*s*_ is the sample refractive index and NA is the numerical aperture of the microscope objective. The location of the foci in each sub-aperture (perspective view), denoted (*x*_*uv*_, *y*_*uv*_) is related to the 3D emitter position (*x*_*i*_, *y*_*i*_, *z*_*i*_) according to:

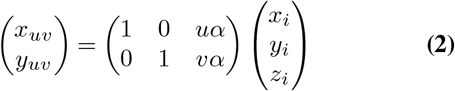

where *α* = *α*(*u, v*) is defined:

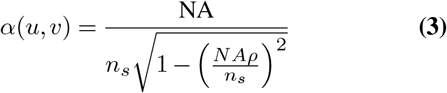

Given a sufficient number of photons, the centre of each foci can be estimated with a precision much finer than it’s width by fitting a 2D Gaussian profile (29–31). An extremely convenient feature of SMLFM is that existing algorithms and software packages (22) designed and optimized for traditional 2D SMLM can be applied to raw SMLFM data to yield a set of *n* localizations {(*x*_*uv*_, *y*_*uv*_)}. ThunderSTORM was used throughout this work (21). Given this set of localizations, the 3D position of a point emitter can be estimated as the least-squares solution to an equation of the form *A***x** = **b**. Here **b** represents the set of 2D localizations, *A* describes the disparity between perspective views as per Equation 2 and **x** = (*x*_*i*_, *y*_*i*_, *z*_*i*_) is the 3D SMLFM localization.

In most SMLM experiments, it is necessary to detect several hundred thousand localizations to generate high resolution datasets and achieve Nyquist sampling of the underlying structure (32). Hence any viable 3D approach must be capable of detecting and localising multiple emitters in each frame. As demonstrated in (c) of Figure 2, in SMLFM this is achieved by using Equations 2 and 3 to identify the most-likely subset of 2D localizations in {(*x*_*uv*_, *y*_*uv*_)} which correspond to a single emitter. Briefly, the set of localizations {(*x*_*uv*_, *y*_*uv*_)} is ordered by decreasing photon number and increasing radial co-ordinate. Taking each member of this ordered set as a ‘seed’ localization, the number of corresponding 2D localizations in {(*x*_*uv*_, *y*_*uv*_)} found at each possible diffraction limited location across the entire axial range is integrated. The largest, and hence most-likely, grouping is identified, and an ordinary least squares solution is calculated to *A***x** = **b**, to yield **x**_*i*_ = (*x*_*i*_, *y*_*i*_, *z*_*i*_). A successful SMLFM localization results in the group of localizations being removed from the available pool. The process is repeated until no more localizations can be grouped and fitted.

Due to sample and system aberrations, the phase in the pupil plane cannot be entirely accounted for by point source displacements. In other 3D SMLM approaches, it is necessary to estimate or calculate Φ_exp_. in order to scale a depth-dependent calibration curve and correct the estimated *z*_*i*_ positions. Phase retrieval methods require stacks of images containing multiple emitters imaged at different depths to calculate the experimental phase (12, 13). This is necessary because in most imaging modalities, angular information is lost when an intensity measurement is made using a detector. Since both intensity and angular information are captured in light field measurements, it is possible to directly measure aberrations, using the 2D localizations themselves, similarly to the method used in (33). For all experimental data presented in this work, these aberrations are estimated by measuring the average residual disparity across the field of view, for emitters close to the focal plane. The residual disparity is subsequently subtracted from all localizations and the light field fitting algorithm re-run to recover the 3D position of point sources. The 3D SMLFM fitting procedure is summarized in Figure 2. For further details, along with a summary of the parameters used for 2D and light field fitting, refer to the supplementary information.

## 3 SMLFM optical design

A standard widefield microscope can be converted to a Fourier light field microscope by adding two components, a lens (*L*_3_) and a microlens array. L_3_ is placed in a 4*f* configuration with the tube lens, *L*_2_, which relays the back focal (Fourier) plane onto the MLA. As is the case with Shack-Hartmann sensors, the performance of SMLFM is primarily dictated by the properties of the microlenses spanning the pupil. The effective pitch determines the extent of the wavefront sampled by a microlens and, further, the division of collected photons into separate foci. Precise light field localization requires finer wavefront sampling which can be achieved using smaller microlenses. Decreasing the microlens pitch increases the axial range since the effective NA of each microlens dictates the operable depth of field. However, the division of photons amongst a large number of microlenses leads to degradation of 2D localization precision.

To investigate these relationships experimentally, two different configurations of light field microscope were built and tested (hereafter referred to as configuration 1 and configuration 2). Since the relative robustness of SMLFM to aberrations reduces the requirement for refractive index matching, an oil immersion lens was used to maximize collection efficiency. The lens used also had a pupil diameter which was easily magnified by the tube lens and off-the-shelf achromatic lenses *L*_3_ = 75 mm (configuration 1) and L_3_ = 100 mm (configuration 2) to a diameter approximately equal to an integer number of microlenses for both configurations. The same, square lattice MLA (SUSS micro-optics, 18-00178) with microlens pitch of 1015 *μ*m and focal length of 25.4 mm was used in both configurations. These two configurations have differing number of illuminated microlenses and magnification factors to the sCMOS sensor located at the focal plane of the MLA. For precise details of both configurations, refer to the supplementary information where further information regarding the consequences of utilizing an oil immersion objective to image into aqueous medium may also be found.

The square lattice of the MLA used in our experiments results in partially illuminated microlenses, as illustrated in (A) of Figures 1 and 3. Since this results in distorted PSF’s all data from these microlenses was excluded from analysis in this proof-of-principle work. As a result, the maximum photon throughput was reduced from a theoretical throughput of 100% to maximum values of 65% (configuration 1) and 87% (configuration 2). The actual throughput depends on the number of views used to estimate emitter position. A custom-designed MLA could be fabricated to optimally tessellate the pupil and maximize the photon throughput.

**Fig. 1.**
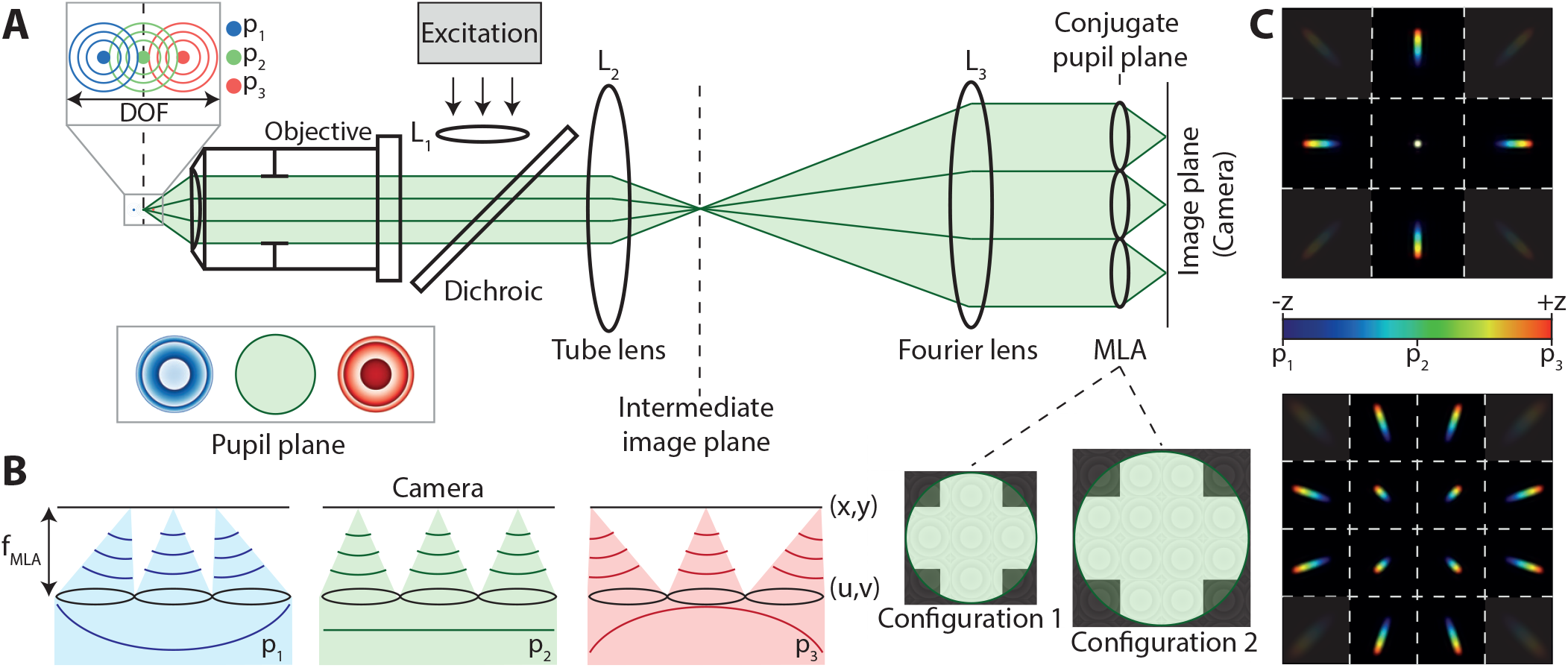
(A) Optical layout of a light field microscope with a microlens array positioned in a conjugate pupil plane. (B) The microlens array samples spatial and angular information from the wavefront, which exhibits asymmetric curvature about the primary image plane. Hence two emitters located at (*x*_*i*_, *y*_*i*_, *z*_*i*_) (red) and (*x*_*i*_, *y*_*i*_, −*z*_*i*_) (blue) are imaged to different positions in each perspective view. (C) Simulated point spread functions for two different light field microscope configurations, with different magnifications of the back focal plane and hence different effective microlens NA (depth of field).

**Fig. 2.**
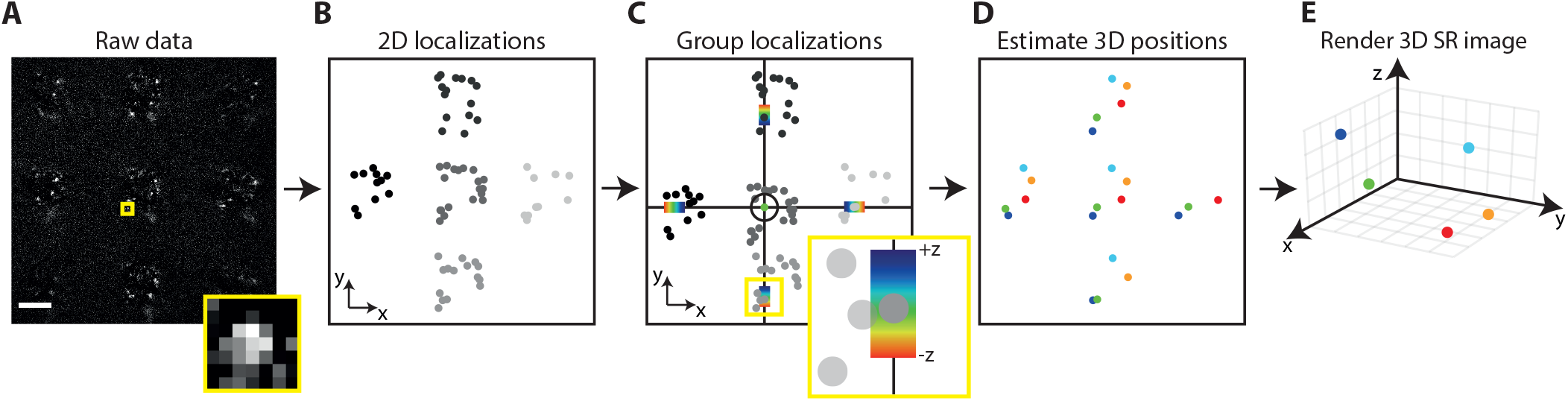
A schematic summary of the algorithm used to estimate 3D emitter position from 4D light field measurements. (A-B) Images of point emitters are detected and localized by Gaussian fitting using traditional 2D SMLM algorithms. Each localization is indexed by the view it appeared in (illustrated here by different shades of grey). Scale bar in (A) represents 15 *μ*m. (A) (inset) Example of an image of a single molecule in a perspective view. (C) localizations in different views corresponding to the same emitter are identified by applying the constraints of the optical model and removed from the set of all 2D localizations. The process is iterated over until there are no more un-grouped 2D localizations (D) The normal least-squares solution is calculated to give each 3D localization. (E) The 3D localizations are plotted to yield a super-resolved image.

**Fig. 3.**
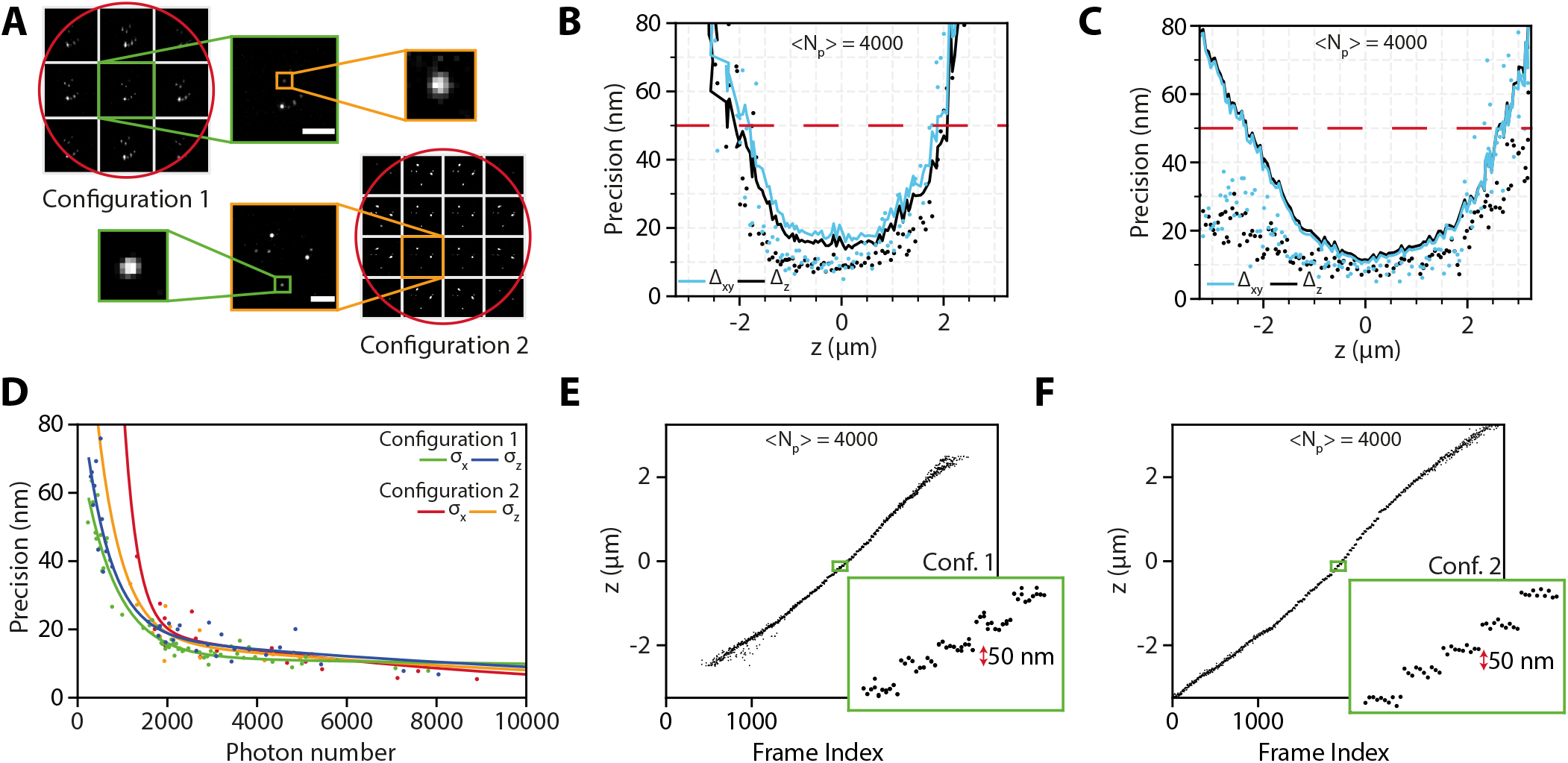
(A) Images of fluorescent beads in different views for configuration 1 and configuration 2. The red circles illustrate the wavefront diameter in the BFP. The white lines indicate the microlens edges. In both configurations, the corner microlenses were partially illuminated and were not included in subsequent analysis. Scale bars represent 1 *μ*m. (B-C) Lateral and axial localization precision (circular markers) and fit error (Δ_*x*_, Δ_*z*_) as a function of the axial position (*z*) of an emitter for (B) configuration 1 and (C) configuration 2. (D) Lateral and axial precision (markers) as a function of number of photons. (E-F) 50 nm axial steps can be resolved using both configurations.

## 4 Results and discussion

To benchmark the performance of SMLFM, a 2D sample comprised of 100 nm fluorescent beads (TetraSpeck Fluorescent Microspheres Kit; T14792; ThermoFisher) immobilized on a coverslip (#1.5 thickness) was imaged. Data was acquired as the microscope stage was translated in fixed steps of 50 nm along the optical axis. For both configurations, 4000 net photons were detected on average across the entire axial range. The localization precision was calculated as the standard deviation of the fitted 3 D p osition a t e ach 50 nm step with 10 repeats at an exposure of 10 ms. A summary of results is presented in in Figure 3. For configuration 1, isotropic lateral and axial localization precision was measured, remaining below 20 nm throughout a 3 *μ*m imaging depth (below 50 nm over a 4 *μ*m axial range). As expected, due to the lower effective NA of each microlens, configuration 2 exhibited a larger depth of field w ith t he isotropic lateral and axial precision remaining below 20 nm over an extended 5 *μ*m range. At photon flux o f 4 000 p er event, this value is competitive with other 3D localization techniques (9, 10, 34). The data presented in (B-C) of Figure 3, demonstrate that the fit error is a robust upper-bound for the precision across all depths and hence can be used to evaluate the quality of each fit.

The relationship between SMLFM localization precision and number of photons was measured by varying the laser power to explore a range of net detected photons. The 10 ms exposure time was kept constant. This data was acquired over an axial range of 4 *μ*m (configuration 1) and 7 *μ*m (configuration 2). The localization precision was again calculated as the standard deviation of the fitted 3D position across 20 repeats (10 repeats for configuration 2). A summary of results is presented in (D) of Figure 3. For the full dataset refer to the supplementary information. At low photon numbers configuration 1 o utperforms c onfiguration 2, du e to th e higher number of photons per microlens, resulting in higher signal-to-noise ratio and better 2D localization precision. However, at sufficiently high photon numbers, the performances of the two configurations becomes comparable. The precision floor of both configurations is approximately isotropic with values of 8 nm (configuration 1) and 10 nm (configuration 2) respectively. A linear, monotonic relationship between fit and stage position was observed in the case of both configuration 1 and configuration 2, demonstrating a 1:1 mapping between emitter location and disparity between perspective views (E-F in Figure 3). Crucially, clear contrast can be observed in the 50 nm axial steps. Taken together, the experimental results presented in Figure 3 confirm the viability of LFM as a single molecule imaging technique.

The single-molecule sensitivity of SMLFM was conclusively demonstrated by imaging Alexa-647 dispersed on a coverslip (#1.5 thickness) with a 70 ms exposure time. Fluorescent traces exhibiting discrete signal levels, characteristic of single molecule photobleaching events were observed (35). Figure 4 shows an example of single step photobleaching of a fluorophore at the tail-end of the distribution of typical localised molecules acquired in this experiment. Traces of the integrated intensity from images of the emitter in each perspective view are plotted. As expected, spatio-temporal correlations are observed between measurements from different views.

**Fig. 4.**
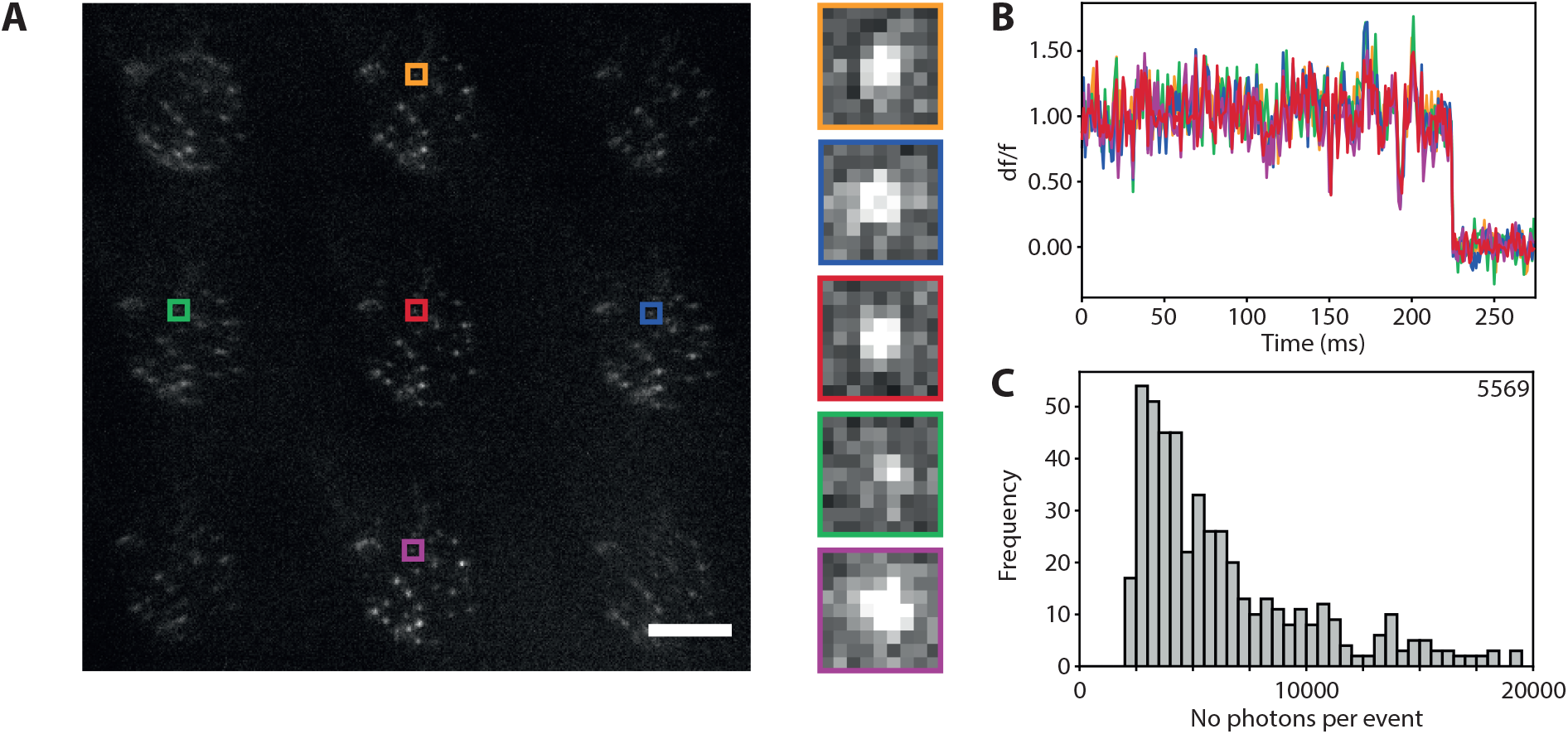
(A) Background subtracted camera frame of Alexa-647 captured using SMLFM (configuration 1). Scale bar represents 15 *μ*m. Insets: images captured in each perspective view. (B) Photobleaching curves of Alexa-647 imaged with light field microscopy. The normalized, integrated intensity in each perspective view is plotted as a function of time. Traces from different perspective views are distinguished by color. (C) Histogram showing the distribution of the number of photons emitted per event. The median number of photons collected per event was 5569.

To examine the super-resolution structural imaging capabilities of SMLFM, we imaged the membrane of fixed Jurkat T-cells using point accumulation for imaging of nanoscale topography (PAINT), based on the stochastic binding of fluorescent wheat germ agglutinin (36). Cells were imaged by using a HILO illumination to reduce background. Datasets comprised of 45, 000 to 150, 000 images were acquired over 1 to 3 hours. For full details of the experimental parameters refer to the supplementary information. Typical frames, which capture information throughout the depth of field, contained 80 2D localizations, corresponding to, on average, 10 total light field localizations. After filtering by precision (using an upper limit of 80 nm) experiments achieved averages between 3 and 13 light field localizations per frame. The density of localizations permitted achieved using SMLFM is ≈ 1.3 million localizations (≈ 380, 000 post-filtering) over 45,000 frames. This high density is enabled by the ability to fit multiple molecules with the same lateral position but differing axial position. This localization density is competitive with other large depth-of-field 3D SMLM approaches (8, 9, 15). Visualizations of these filtered SMLFM localizations are shown in Figure 5 for two such experiments, one for each SMLFM configuration. Horizontal and vertical projections through each cell demonstrate that the resolution of SMLFM is sufficient to resolve the 3D membrane contour and microvilli.

**Fig. 5.**
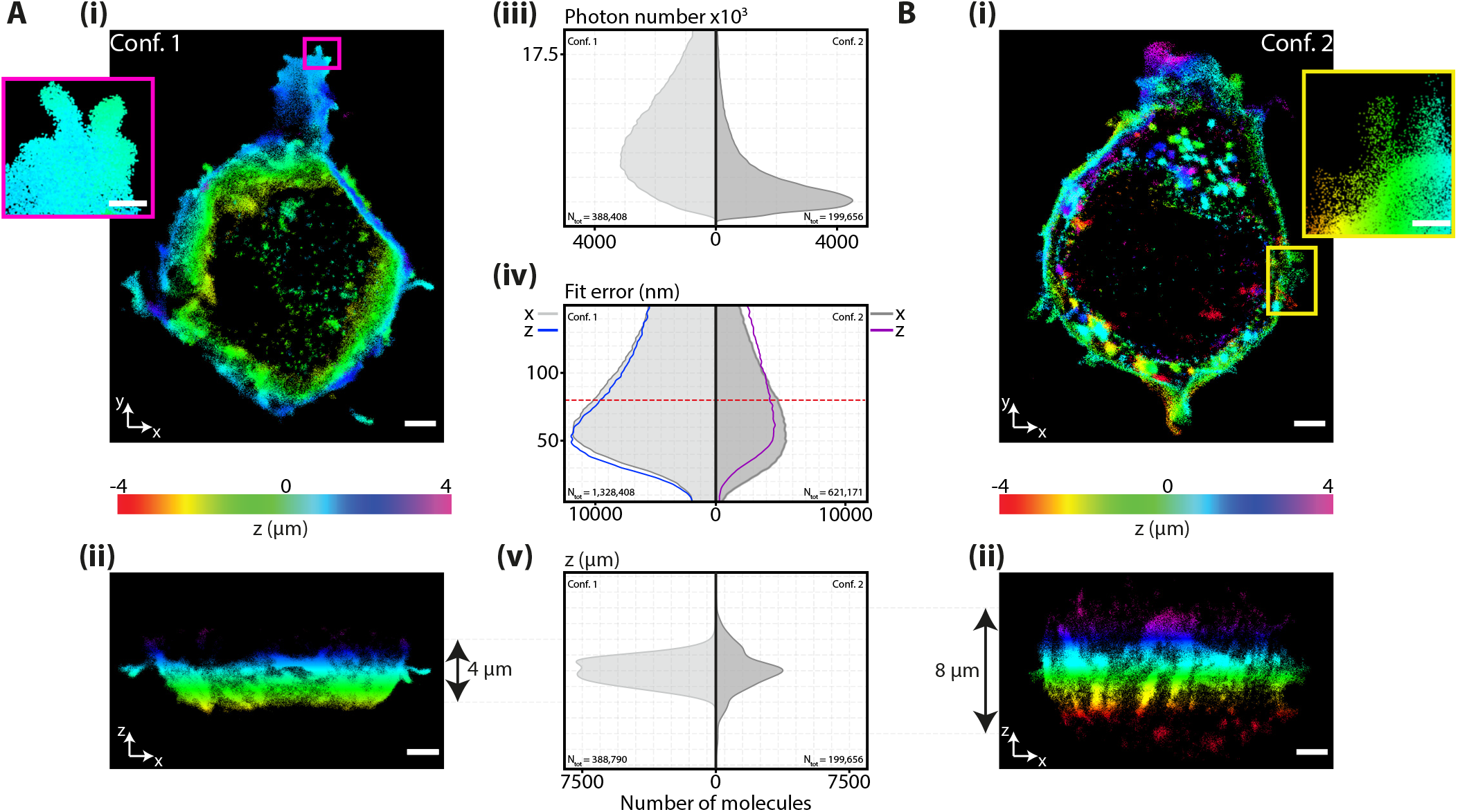
Super resolved images of Jurkat T cells captured with Single Molecule Light Field Microscopy. (A) (i) Horizontal and vertical (ii) cross-sections through data acquired with configuration 1. Scale bar represents 2 *μ*m. (i) Inset: zoomed image of microvilli. Scale bar represents 500 nm. (A) (iii - v) kernel density plots summarizing the characteristics of the acquired data. (iii) and (v) include data filtered into the final visualizations. The fit error for all light field localizations is plotted in (iv). Data with fit error below 80 nm, indicated by the red dashed line was filtered into the final visualizations. (B) (i - v) as for (A) for data acquired with configuration 2.

## 5 Conclusion

We have demonstrated the viability of SMLFM for scanless 3D super-resolution imaging. Our results show that SMLFM can localize single molecules with a near isotropic precision of 20 nm using only a few thousand emitted photons, a comparable performance to other 3D imaging techniques (3–6). We have also demonstrated detection and 3D localization of single molecules in densely blinking specimens, achieving up to 25 light field localizations per frame in data sets of 40, 000 to 150, 000 frames. The mechanism which enables SMLFM, disparity between perspective views, is one which reveals the underlying wavefront structure and amplitude of the field in the pupil. Such data enables post-acquisition aberration correction without requiring phase retrieval or z-dependent calibration scans. This rich information coupled with the simple PSF footprint and the optical properties of microlens arrays result in SMLFM having the potential to offer highly accurate and precise multi-colour 3D nanoscopy over whole eukaryotic cell volumes.

See Supplement 1 for supporting content.

## Author contributions

KOH conceptualized the project. KOH and SFL supervised and administered the project. RRS, SAR, LAM and KOH developed the methodology. RRS, SAR, AJC, MOL and KOH conducted the experimental investigation. ES and AP provided the cell samples and labelling methodology. RRS, SAR, LAM and KOH performed formal analysis of data and developed software. RRS was responsible for visualization of results in the manuscript. RRS, SAR and KOH wrote the manuscript. All authors reviewed and edited the final manuscript.

## ACKNOWLEDGEMENTS

This work was funded by the EPSRC (EP/L015455/1, EP/R025398/1) and The Royal Society University Research Fellowship to SFL (URF/R/80029).

