## Supplementary material for "Single Molecule Light Field Microscopy"

### Single Molecule Light Field Microscopy: supplementary material

#### 1 Estimation of 3D emitter position from SMLFM measurements

In an ideal microscope, a spherical wave emitted by a point source located at the origin $(x_p, y_p, z_p)$ is transformed into a plane wave at the pupil. Spherical waves emitted by a point emitter displaced from the origin, located at $(x_i, y_i, z_i)$, generate more complex wavefronts in the pupil since the optical path lengths to the principal sphere differ for rays propagating at different angles (spatial frequencies). The majority of microscope objectives are designed according to the sine condition, according to which, ray height is preserved during mapping between principal planes $(\{x_p, y_p\} \rightarrow \{u, v\})$ (11). As a result, the phase difference in the pupil between wavefronts emanating from points located at $(x_p, y_p, z_p)$ and $(x_i, y_i, z_i)$ depends on the optical path difference (OPD) between two rays that intersect at the principal sphere, as depicted in Figure 1:

$$\text{OPD} = n_s\left(|\vec{d}| - f\right) = n_s\left(|\vec{f} - \vec{r}| - f\right)$$

$$= n_s\sqrt{(x_p - x_i)^2 + (y_p - y_i)^2 + \left(\sqrt{f^2 - x_p^2 - y_p^2} - z_i\right)^2} - n_s f \quad \textbf{(1)}$$

For small displacements from the focal point, the quotient $\left(x_i^2 + y_i^2 + z_i^2\right)/f^2$ vanishes. To first order expansion, Equation 1 takes the form:

$$\text{OPD} \sim \frac{n_s}{f}\left[(x_i x_p + y_i y_p) + z_i\sqrt{f^2 - x_p^2 - y_p^2}\right] \quad \textbf{(2)}$$

Converting OPD to phase by multiplying by the free space wavenumber $k = 2\pi/\lambda$, the phase in the pupil plane resulting from displacements from the focal position is:

$$\Phi_{x_i, y_i, z_i} = \frac{n_s k}{f}\left[(x_i u + y_i v) + z_i\sqrt{f^2 - u^2 - v^2}\right] \quad \textbf{(3)}$$

In normalized pupil co-ordinates, where $\rho^2 = u^2 + v^2 = 1$ at the pupil edge and 0 at the optical axis, Equation 3 becomes:

$$\Phi_{x_i, y_i, z_i} = k\text{NA}\,(x_i u + y_i v) + n_s z_i k\sqrt{1 - \left(\frac{\text{NA}\rho}{n_s}\right)^2} \quad \textbf{(4)}$$

In Fourier light field microscopy, as in Shack-Hartmann wavefront sensors, $\Phi$ is sampled by a microlens array. Each microlens generates a focused image, displaced in the direction of and at a distance proportional to the local phase gradient $(\partial\Phi/\partial\rho)$, averaged across each sub-aperture. The location of the foci in each perspective view (in sample co-ordinates), denoted $(x_{uv}, y_{uv})$ is given:

$$\begin{pmatrix} x_{uv} \\ y_{uv} \end{pmatrix} = \frac{1}{k\text{NA}}\begin{pmatrix} \frac{\partial\Phi_u}{\partial u} & \frac{\partial\Phi_v}{\partial u} \\ \frac{\partial\Phi_u}{\partial v} & \frac{\partial\Phi_v}{\partial v} \end{pmatrix} \quad \textbf{(5)}$$

where the factor $1/\text{NA}$ accounts for the transformation between image and object space (or, equivalently, pupil plane co-ordinates to object space co-ordinates). In the absence of aberrations, the phase in the pupil plane is entirely due to point source displacements from the focal position. Hence the derivative of the phase along each principal axis is given by:

$$\frac{\partial\Phi_u}{\partial u} = k\text{NA}x_i - \frac{k\text{NA}^2 u z_i}{n_s\sqrt{1 - \left(\frac{\text{NA}\rho}{n_s}\right)^2}} \quad \textbf{(6)}$$

Such that Equation 5 may be written in the form:

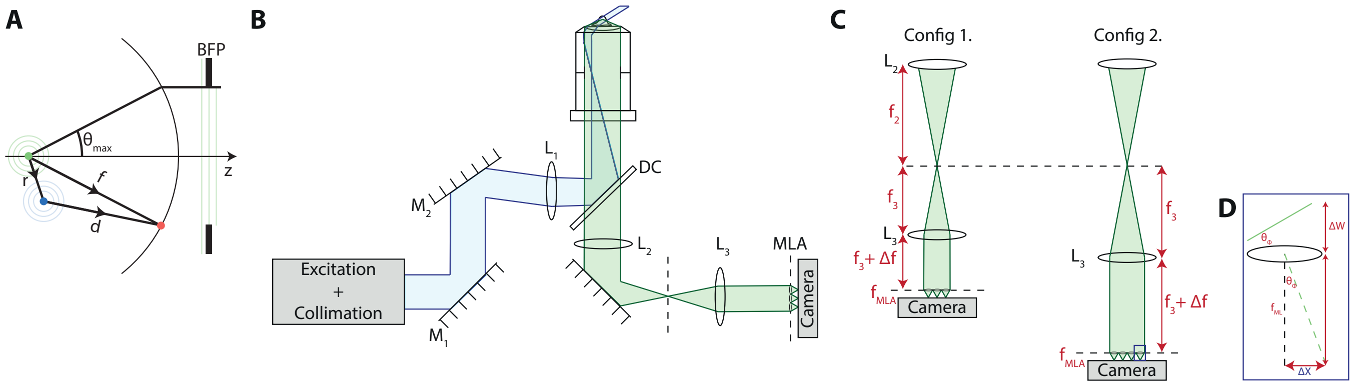


**Fig. 1.** (A) Geometry for calculating the optical path difference between wavefronts from an emitter (green) located at the focal point, $(x_p, y_p, z_p)$, and an emitter displaced from the focal point (blue) to $(x_i, y_i, z_i)$ which intersect at a point on the principal sphere (red). (B) Schematic diagram of experimental setup. In the excitation path (blue) a collimated laser beam is focused to the edge of the back aperture of the objective for HILO illumination of the sample. In the detection path (green), the back focal plane is magnified with lenses $L_2$ and $L_3$ positioned in a 4f configuration. The microlens array is positioned in the conjugate pupil plane. (C) The size of the back focal plane is dictated by the focal length of $L_3$. This changes the number of microlenses which span the magnified pupil. (D) The position of the image behind each microlens (here denoted $\Delta X$) is proportional to the average tilt of the wavefront spanning the diameter $\Delta W$.

$$\begin{pmatrix} x_{uv} \\ y_{uv} \end{pmatrix} = \begin{pmatrix} 1 & 0 & u\alpha \\ 0 & 1 & v\alpha \end{pmatrix} \begin{pmatrix} x_i \\ y_i \\ z_i \end{pmatrix} \quad \textbf{(7)}$$

where $\alpha(u, v)$ is defined:

$$\alpha(u, v) = \frac{-\text{NA}}{n_s \sqrt{1 - \left(\frac{NA\rho}{n_s}\right)^2}} \quad \textbf{(8)}$$

This Equation corresponds to Equation (3) in the main text. Note that the parity of $\alpha(u, v)$ depends on the convention used to describe positive and negative axial displacements from the focal plane.

In SMLFM, a 2D Gaussian profile is fit to the image of every point emitter in each perspective view (the sensor pixels bounded by each microlens). Given this set of localisations, the 3D position of a point emitter can be estimated as the least-squares solution to $A\mathbf{x} = \mathbf{b}$. Here $\mathbf{b}$ represents the 2D localisations, $A$ describes the disparity between perspective views as per Equation **??** and $\mathbf{x}_i = (x_i, y_i, z_i)$ is the 3D SMLFM localisation. Written explicitly:

$$\begin{bmatrix} 1 & 0 & u_1\alpha(u_1, v_1) \\ \vdots & \vdots & \vdots \\ 1 & 0 & u_n\alpha(u_n, v_n) \\ 0 & 1 & v_1\alpha(u_1, v_1) \\ \vdots & \vdots & \vdots \\ 0 & 1 & v_n\alpha(u_n, v_n) \end{bmatrix} \begin{bmatrix} x_i \\ y_i \\ z_i \end{bmatrix} = \begin{bmatrix} x_1 \\ \vdots \\ x_n \\ y_1 \\ \vdots \\ y_n \end{bmatrix} \quad \textbf{(9)}$$

#### 2 SMLFM and Aberrations

In the prototype system presented in this work, an oil immersion objective was used to image T-cells mounted in an aqueous medium ($n_s \approx 1.33$). The oil immersion lens was used as a result of the convenient size of its back focal plane, which was magnified using off-the-shelf achromatic lenses to a diameter approximately equal to an integer number of microlenses for both configurations. Oil immersion objectives are generally designed for stigmatic imaging of points located on the 'design surface', which typically lies immediately below the cover slip surface. Furthermore, stigmatic imaging only occurs when a certain thickness of immersion oil and cover slip are used. The results presented in Figure 4 in the main text were collected under experimental conditions depicted in Figure 2. Principally, emitters were located below a $\#1.5$ coverslip, immersed in a medium with refractive index $n_s$, and imaged with an objective lens designed for an immersion medium with refractive index $n_{\text{imm.}}$. In the ideal microscope described in Section 1, the phase in the pupil plane was considered entirely due to point source displacements from the focal position, described by Equation 4. According to this model, laterally displaced point sources on the nominal focal plane are imaged to identical locations in each perspective view in SMLFM. This is not the case when imaging into an aqueous medium using an oil immersion objective, a well characterized situation (3–6). Following these treatments, consider a point source located at $(x_i, y_i, z_i) = (0, 0, 0)$, a distance $d$ below the cover slip. Figure 2 demonstrates how refraction of rays at the sample-cover slip interface results in a so-called 'focal-shift': the closest approximation to

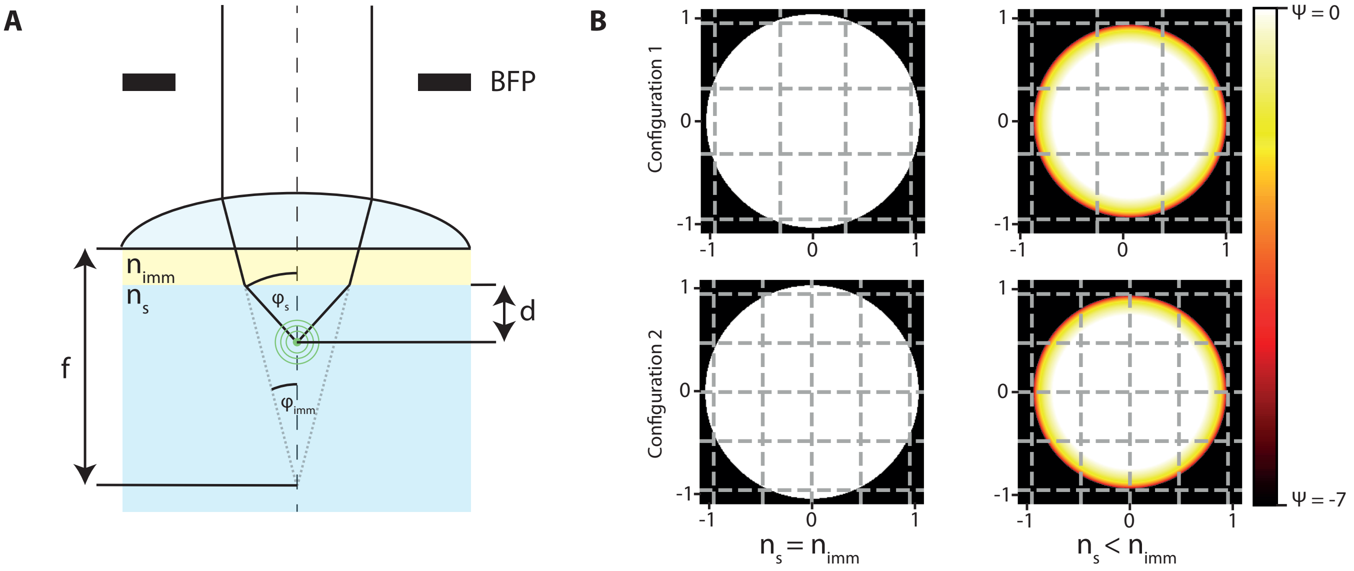


**Fig. 2.** (A) Imaging into aqueous medium ($n_s < n_{\text{imm.}}$) using an oil immersion objective. Refraction at the boundary between the cover slip and the aqueous medium results in a focal shift. (B) For $\varphi_{\text{imm.}} > \theta_{\text{crit.}}$ the amplitude of the electric field in the pupil plane is zero for emitters located $d > \lambda_{\text{em.}}$ away from the cover slip. Configurations 1 and 2 were designed such that an integer number of microlenses (depicted by gray dashed lines) span this effective reduced pupil diameter.

stigmatic imaging occurs at a plane located a distance $d$ below the coverslip when the $\rho^2$ terms of spherical aberration and defocus compensate one another.

Based on the assumptions that the surface of the front lens of the microscope objective is planar, and also that the microscope objective satisfies the sine condition, the phase in the back focal plane due to this focal-shift (or, spherical aberration) can be calculated by calculating the optical path difference $\text{OPD}_{\text{sph.}}$ between rays in the design system and in the experimental system which enter the the microscope objective as parallel rays:

$$\text{OPD}_{\text{sph.}} = d\left(n_s \cos\varphi_2 - n_{\text{imm.}} \cos\varphi_1\right) \tag{10}$$

where, according to the sine condition:

$$\sin\varphi_1 = \frac{\text{NA}\rho}{n_{\text{imm.}}} \tag{11}$$

and, from Snell's law:

$$\sin\varphi_2 = \frac{\text{NA}\rho}{n_{\text{s}}} \tag{12}$$

such that the full expression for the phase in the pupil is:

$$\Phi_{\text{sph.}} = kd\left[n_s \cos\left(\arcsin\left(\frac{\text{NA}\rho}{n_s}\right)\right) - n_{\text{imm.}} \cos\left(\arcsin\left(\frac{\text{NA}\rho}{n_{\text{imm.}}}\right)\right)\right] \tag{13}$$

with corresponding $\alpha_{\text{sph.}}$:

$$\alpha_{\text{sph.}} = 2d\left[\frac{\text{NA}^2\rho^2}{n_{\text{imm.}}\sqrt{1-\left(\frac{\text{NA}\rho}{n_{\text{imm.}}}\right)^2}} - \frac{\text{NA}^2\rho^2}{n_{\text{s}}\sqrt{1-\left(\frac{\text{NA}\rho}{n_{\text{s}}}\right)^2}}\right] \tag{14}$$

As a result of spherical aberration, even a point emitter located at the nominal focal plane will be imaged to different locations in different perspective views in SMLFM. This is demonstrated in Figure 2, which depicts the phase in the back focal plane due to an emitter at the nominal focal plane. The total phase for emitters located at $(x_i, y_i, z_i)$ is now the sum of two terms:

$$\Phi_{\text{tot.}} = \Phi_{x_i,y_i,z_i} + \Phi_{\text{sph.}} \tag{15}$$

In other 3D SMLM approaches, it is necessary to estimate or calculate $\Phi_{\text{sph.}}$ in order to scale a calibration curve and correct the estimated $z_i$ positions. One such approach is phase retrieval, which requires a z stack of a bead on a cover slip to calculate the experimental phase (1, 2). This is necessary because in most imaging modalities, angular information is lost when an intensity measurement is made using a detector. As described in the main text, in SMLFM, both intensity and angular information is captured. As a result it is possible to directly measure spherical aberration, and other aberrations, using the 2D localisations themselves. For all experimental data, these aberrations are estimated by measuring the average residual disparity across the field of view $< \alpha(u,v)|_{z=0} >$, for emitters located at the focal plane. The residual disparity is subsequently subtracted from all frames.

There are other subtleties to be considered when using an oil immersion lens to image into an aqueous medium. Firstly, the maximum solid angle, $\theta_{\text{max}}$ of the microscope objective lens ($\theta_{\text{max}} = 67°$) is larger than the critical angle of refraction ($\theta_{\text{crit.}}$) between the aqueous medium and glass cover slip. As a result, for emitters located greater than a distance $\lambda_{\text{em}}$ away from the cover slip, the amplitude of the electric field above $\rho_{\text{crit.}}$ is zero in the pupil (8–10). Furthermore, refraction at the interface means that each point in the pupil plane receives light from a different solid angle than the case where the sample refractive index matches that of the immersion oil. Furthermore, polarization dependent Fresnel reflections at the sample-cover slip interface reduce the amplitude of the electric field in the pupil plane. Considering these losses is beyond the scope of this work, but would be important for the application of SMLFM to other single molecule experiments.

Other deviations from the ideal case which affect fit quality are misalignments of the microlens array. Lateral misalignments are the least critical since they are equivalent to adding the same tilt to all wavefronts, corresponding to a global translation of all the images in the perspective views. It is however, important to ensure that the outer microlenses along the principal axes are not clipped. Practically speaking this is achieved by minimizing aberrations in the point spread function. It is also necessary to estimate the location of the optical axis relative to the centre of the microlens array. Axial displacements of the sensor from the focal plane decrease precision as photons are distributed over a larger number of pixels due to defocus. Furthermore, a given wavefront tilt will correspond to different $(x_{uv}, y_{uv})$ displacements. Similarly to other wavefront engineering approaches, placing the microlens array in a plane other than a conjugate pupil plane will introduce errors since points from in different areas of the field of view will experience different phase modulation. Rotation of the microlens is not intrinsically problematic, unless it results in clipped microlenses, but the principal axes $(u,v)$ must be identified in order to accurately solve Equation 9. Rotating the set of 2D localisations $\{(x_{uv}, y_{uv})\}$ and maximizing the number of groups of 2D localisations which solved Equation 9 with low fit error as a function of the rotation angle $\theta_{\text{MLA}}$ was found to be a good angular discriminator.

#### 3 Implementation (software)

The procedure used to estimate 3D emitter position from a set of 4D light field measurements may be summarized as:

1. Run 2D SMLM detection and Gaussian fitting. Output: set of localisations $\{(x_{\text{pix}}, y_{\text{pix}})\}$
2. Transform $\{(x_{\text{pix}}, y_{\text{pix}})\} \rightarrow \{(x_{\text{uv}}, y_{\text{uv}})\}$ by translating and scaling
3. Filter $\{(x_{\text{uv}}, y_{\text{uv}})\}$ according to the standard deviation of the Gaussian fit. Remove localisations in external lenses.
4. Estimate $\theta_{\text{MLA}}$ as that which maximizes number of fitted light field points. Rotate $\{(x_{\text{u}}, y_{\text{v}})\}$ by $\theta_{\text{MLA}}$ .
5. Estimate system and sample aberrations ($\langle \alpha(u,v)|_{z=0} \rangle$) as the residual disparity around $z = 0$ within an axial range $\Delta$z. Subtract $\langle \alpha(u,v)|_{z=0} \rangle$ from all localisations $\{(x_{\text{u}}, y_{\text{v}})\}$
6. Order the $q$ localisations in a view close to the optical axis by decreasing number of photons. Take each of the $q$ localisations as a seed and:
   (a) Identify candidate localisations in other views within a permitted disparity range according to the chosen optical model (Equation 8)
   (b) If there are multiple candidates within a given view, take the best fit
   (c) If corresponding localisations found in a minimum of other views, calculate the least squares solution to Equation 9
   (d) If a sufficiently good solution found (residuals smaller than max fit distance), remove these localisations from the pool

For all data presented in the paper, 2D detection and localisation of spots was carried out on raw frames using ThunderSTORM (Fiji plugin) (15). Spots with a standard deviation between $\sigma_{min}$ and $\sigma_{max}$ (see Table 1) were kept for the light field analysis. Localisations from the fully illuminated microlenses were used in the analysis. This resulted in a maximum of 5 (configuration 1) and 12 (configuration 2) 2D fits per emitter for the subsequent light field fit.

Once the seed for the light field fit was selected in a given view, candidate 2D localisations were searched in all the other views. To account for aberrations and the precision of 2D localisations, candidates in other views subtending an angle $\theta_{search}$ with the seed and within a permitted disparity range were considered. Once a minimum number of localizations, $N_{min}$, were found for a given seed, light field fit was performed. If the fit error was below the chosen threshold, the light field fit was selected. A summary of the manual parameters and values found to optimize fit results is presented in Table 1. Whilst this procedure is generally applicable to all SMLFM data, optimization of the parameters in Table 1 is necessary for use with data acquired on systems substantially different than those described in this work.

#### 4 Implementation (hardware)

All Experiments were performed on a customized, inverted microscope (Nikon, Eclipse Ti2). Illumination was provided by a set of lasers (405, 488, 561 and 638 nm) housed in an Omicron LightHUB. The output from each laser was coupled into a single mode optical fibre and collimated into an 9.6 mm diameter beam, using an off-axis parabolic mirror (Thorlabs RC08APC-P01). This beam was focused onto the back focal plane of a $60\times$, 1.42NA oil immersion objective (Olympus, PlanApo N) using a 300 mm focal length lens, $L_1$, to achieve wide-field excitation (96 $\mu$m FWHM). Where specified, data was acquired under Highly Inclined Laminated Optical (HILO) illumination (12), achieved by laterally displacing the excitation beam towards the edge of the back focal plane. A quad-band filter located between $L_1$ and the objective separated the incoming excitation beams from the collected fluorescence. A 200 mm tube lens, $L_2$ focused the collected fluorescence onto an intermediate image plane. To perform SMLFM two more elements are required: another lens, $L_3$, placed in a $4f$ configuration with $L_2$ which relays the back focal (Fourier) plane onto the Microlens Array (MLA). The choice of the focal length of $L_3$ is crucial in determining the depth-of-field and efficiency of SMLFM for a given MLA. We used a square lattice MLA (SUSS micro-optics, 18-00178) with microlens pitch of 1015 $\mu$m and focal length of 25.4 mm. $L_3$ focal lengths of 75 mm and 100 mm were used to generate two different BFP diameter configurations (configuration 1 and configuration 2). These two configurations, illustrated in Figure 1, have differing number of illuminated microlenses and magnification factors to the detector (Hamamatsu, Flash 4.0 v2) which sits at the focal plane of the MLA.

#### 5 Performance

Firstly, the accuracy of the least-squares solution to Equation 9 was evaluated by comparing the average fit error to the standard deviation of location estimates of repeat measurements on multiple images of beads imaged at low laser power. The results are presented in Figure 3 which demonstrates that the fit error is an upper bound for the precision of the estimation of the location of the emitter. Hence the fit error can be used to assess the quality of each fit.

Next, the relationship between localisation precision and number of photons was measured by varying the laser power to explore a range of net detected photons up to $10,000$. Data plotted in Figure 4 demonstrates that the precision floor of both configurations is approximately isotropic with values of 8 nm (configuration 1) and 10 nm respectively.

For point emitters located on the nominal focal plane of the microscope objective $(z_i = 0)$, sharp images are formed in each perspective view. According to Equation 6, the curvature of the wavefront in the pupil increases as a function of $|z_i|$ which leads to defocused images. Since the rate of increase of curvature is fastest at the edge of the pupil, the peripheral microlenses exhibit the lowest depths of field. This is demonstrated in Figure 5 where the average width of the 2D Gaussian fit to an image of a point emitter is plotted as a function of axial emitter position. The asymmetry of curves plotted in Figure 5 indicates that the curvature of the wavefront changes at different rates along the principal as a function of axial position, indicative of an astigmatic aberration. In SMLFM, aberrations in the system are estimated as the residual disparity between localisations corresponding to emitters close to the focal plane. This residual disparity is subtracted from all 2D localisations prior to estimation of 3D emitter location. The fit error for all 3D localisations is plotted in Figure 6 for all datasets presented in this work. In all cases, subtraction of the residual disparity improves the quality of the fit.

| Parameter (units) | Configuration 1 | Configuration 2 |
|---|---|---|
| Max no. of 2D localisations per emitter | 5 | 12 |
| $\sigma_{min}$ for 2D fits (no. pixels) | 0.9 | 0.9 |
| $\sigma_{max}$ for 2D fits (no. pixels) | 3.5 | 3.5 |
| $\theta_{search}$ for candidates ($^\circ$) | 1 | 1 |
| Permitted disparity range (no. pixels) | 0.3 | 0.3 |
| Min. no. of views per LF fit, $N_{min}$ | 2 | 2 |
| 3D Fit threshold ($\mu$m) | 1 | 1 |
| Range for aberration calculation, $\Delta$z ($\mu$m) | 1 | 1 |

**Table 1.** Summary of fit parameters for algorithm. For an explanation of terms, refer to the summary of the algorithm presented in Section 3.

| | Configuration 1 | Configuration 2 | Widefield (2D SMLM) |
|---|---|---|---|
| Magnification | 23.2 | 17.4 | 66.7 |
| Pixel size (nm) | 280.0 | 373.4 | 97.5 |
| DOF ($\mu$m) | 6.36 | 3.57 | 0.36 |
| Max. efficiency (%) | 66 | 87 | 100 |
| $NA_{eff}$ | 0.45 | 0.34 | - |
| Resolution (nm) | 576 | 768.5 | 183 |
| FOV ($\mu$m) | 43.7 | 58.3 | 198.7 |

**Table 2.** Comparison of characteristics of configuration 1 and configuration 2. $NA_{eff}$ refers to the effective numerical aperture of each microlens. Resolution is stated according to the Abbe limit.

| Parameter (units) | Fig. 3B | Fig. 3C | Fig. 3D (conf 1) | Fig. 3D (conf 2) | Fig. 3E | Fig. 3F |
|---|---|---|---|---|---|---|
| $\lambda$ (nm) | 638 | 638 | 638 | 638 | 638 | 638 |
| Avg no. photons | $\sim$ 3800 | $\sim$ 4300 | as plotted | as plotted | $\sim$ 3800 | $\sim$ 4300 |
| Exposure time (ms) | 10 | 10 | 10 | 10 | 10 | 10 |
| No. frames per position | 10 | 10 | 20 | 10 | 10 | 10 |
| Z interval (nm) | 10 | 10 | 100 | 50 | 10 | 10 |
| Z range ($\mu$m) | 4.7 | 6.5 | 4 | 7 | 4.7 | 6.5 |

**Table 3.** Summary of parameters used to acquire data presented in Figure 2 of the main text. The header in each column refers to label of the figure.

| Parameter (units) | Fig. 4A | Fig. 4B |
|---|---|---|
| $\lambda$ (nm) | 638 | 638 |
| Power (kW cm$^{-2}$) | $\sim$ 0.8 | $\sim$ 0.4 |
| Exposure time (ms) | 70 | 50 |
| No. acquired frames | 44676 | 150000 |

**Table 4.** Summary of parameters used to acquire data presented in Figure 4 of the main text. The header in each column refers to label of the figure.

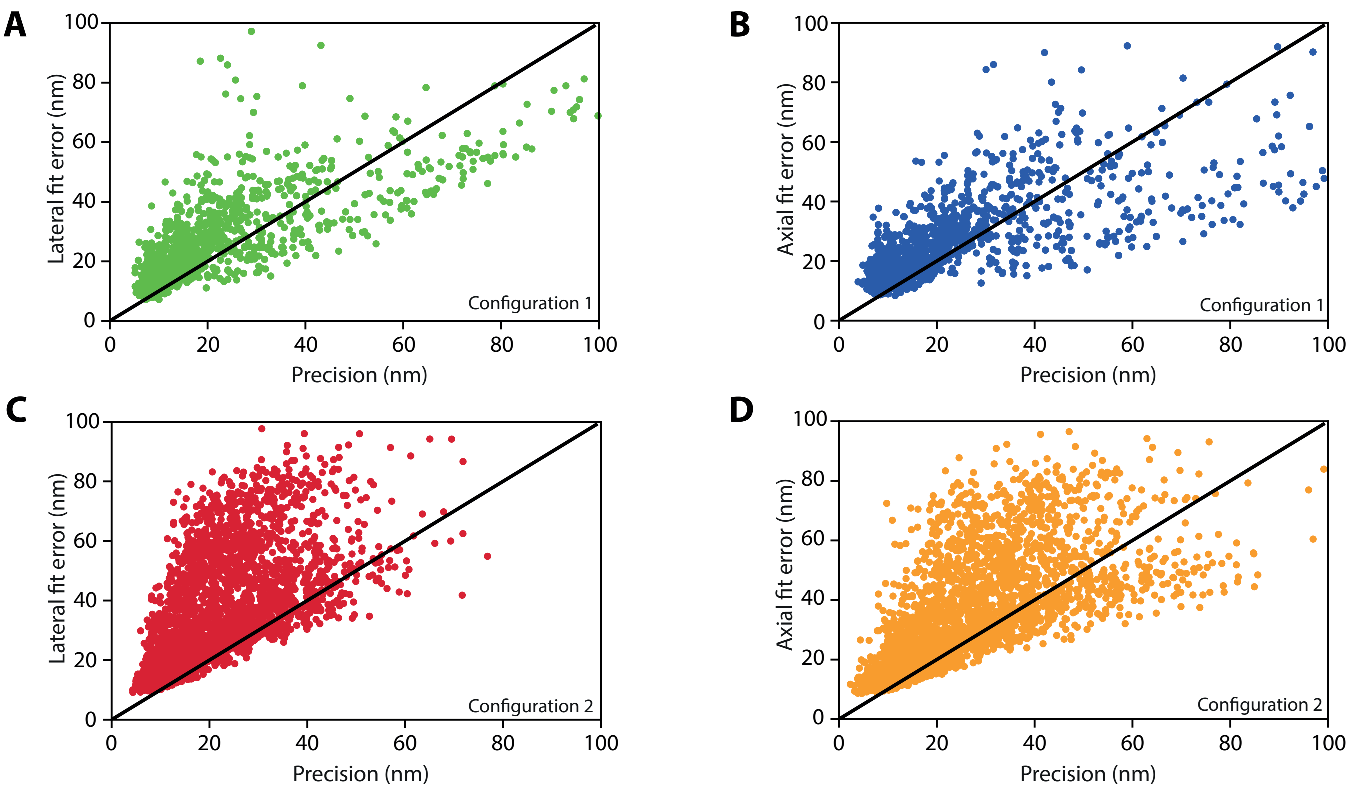


**Fig. 3.** The fit error is an upper bound for the precision of the estimation of the location of the emitter for data acquired in both configurations.

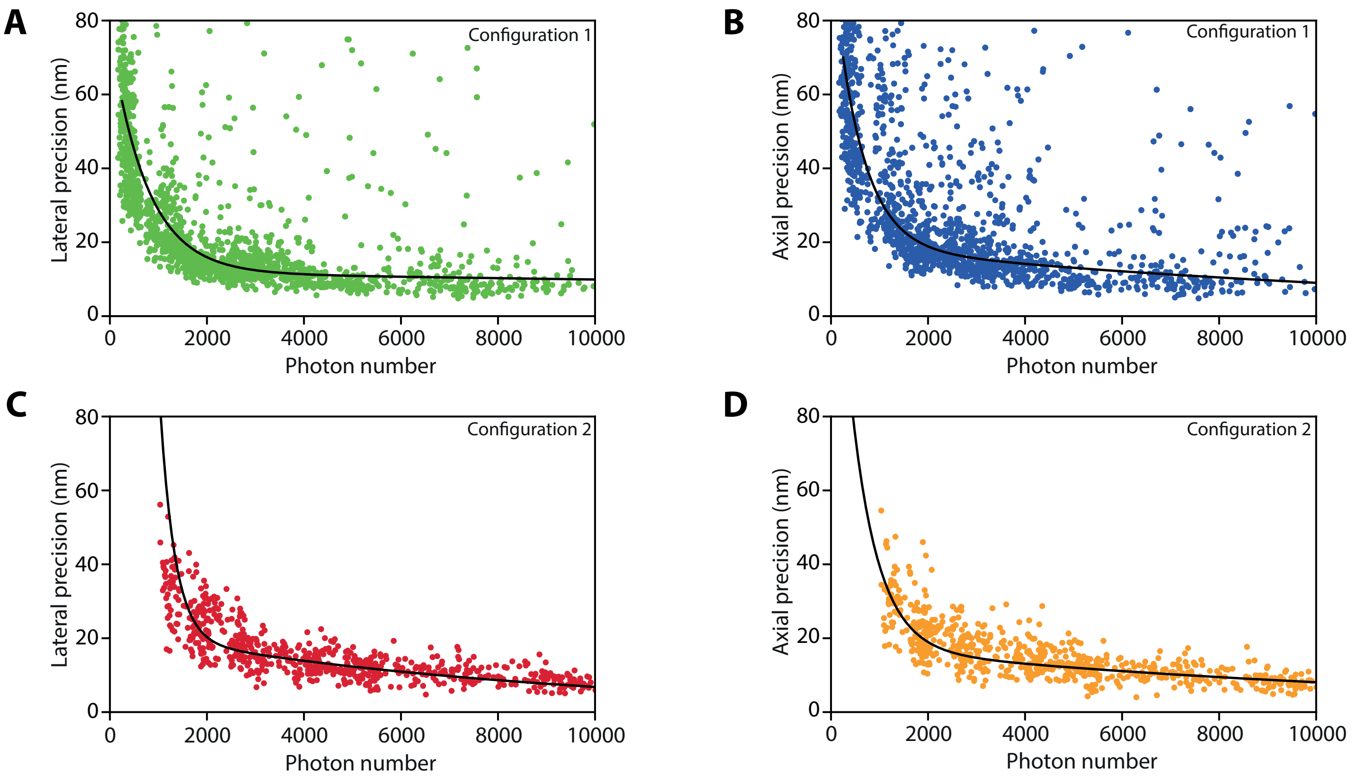


**Fig. 4.** Lateral and axial precision as a function of number of detected photons for each configuration. Top row contains data from configuration 1. Bottom row containst data from configuration 2. In each case a double exponential fit to the data is also plotted (black) to guide the eye.

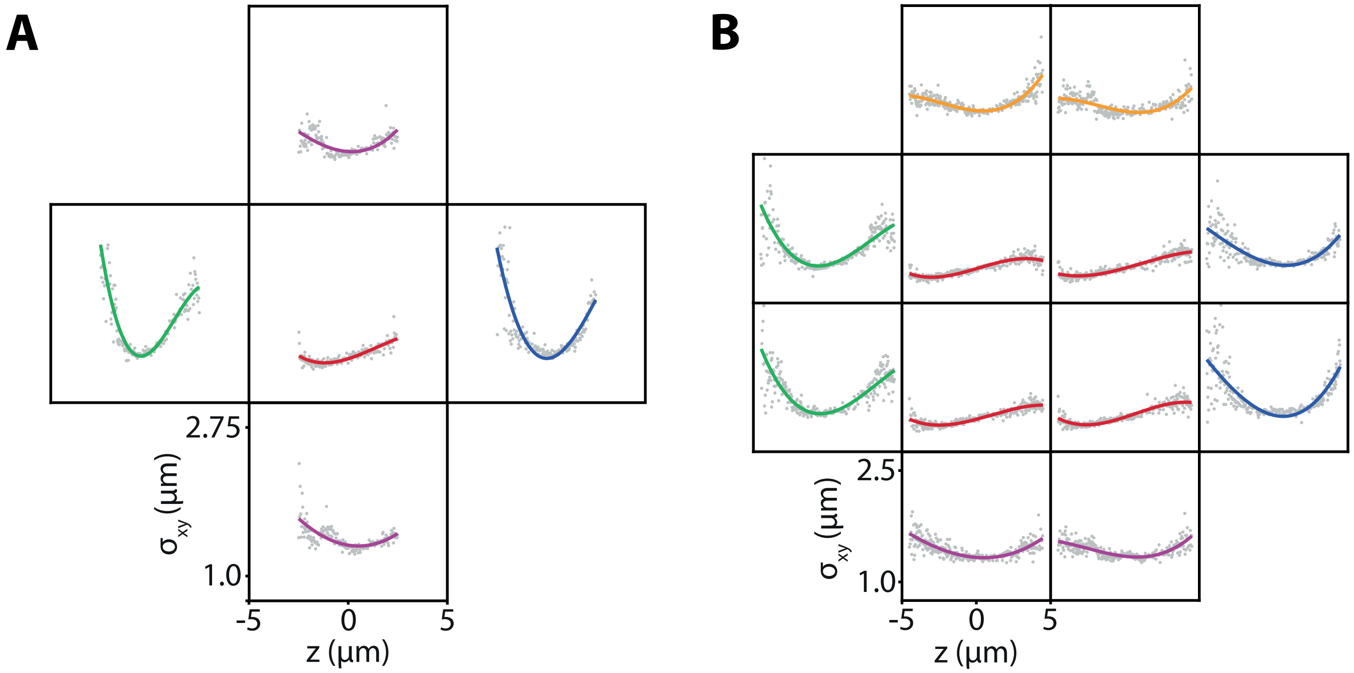


**Fig. 5.** Average width of the 2D Gaussian fit to the image of an emitter in each perspective view as a function of $z$. (A) Configuration 1, (B) Configuration 2. The asymmetry in beam width as a function of axial position clearly present in both cases between the principal axes indicates the presence of astigmatism.

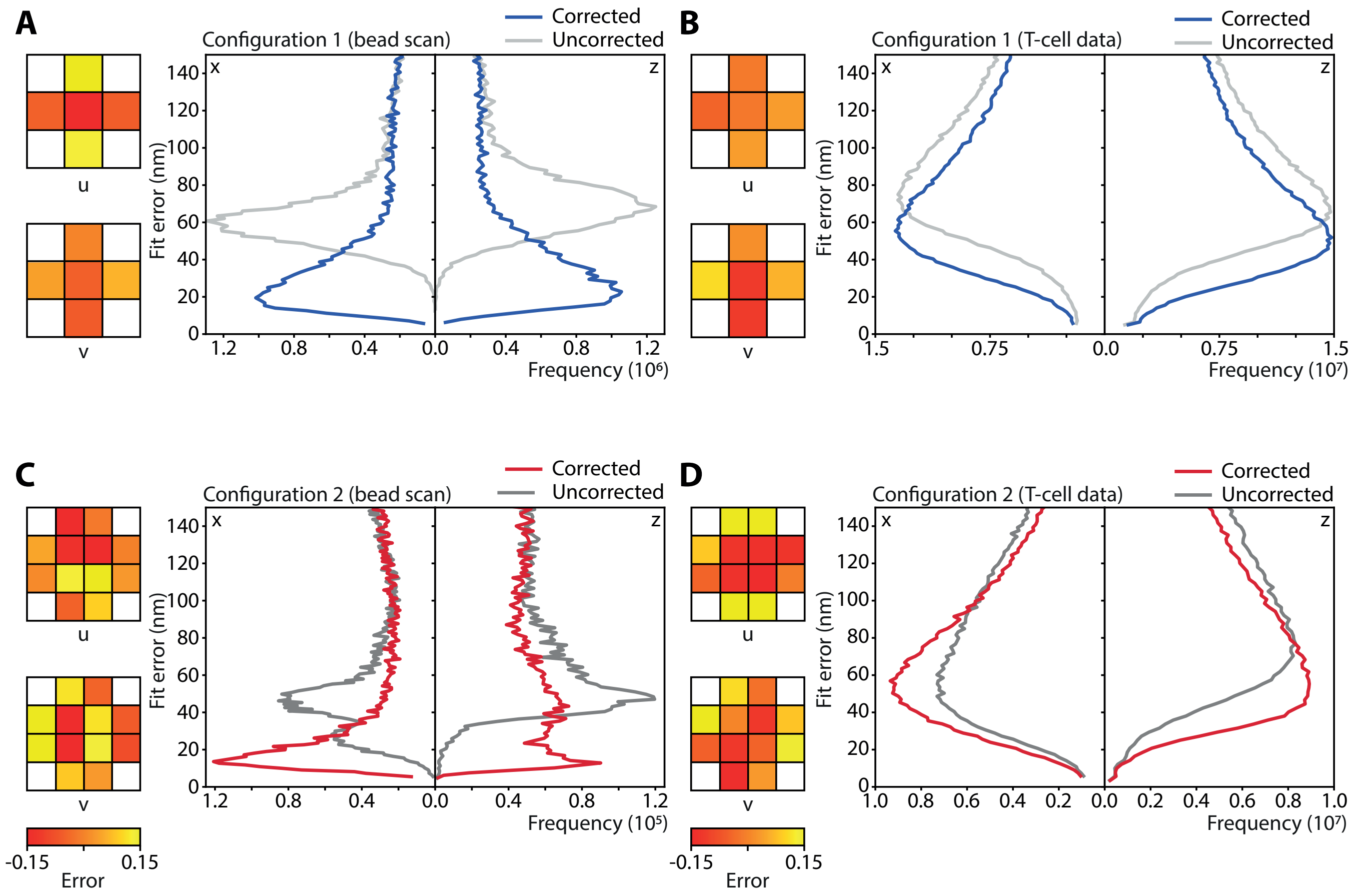


**Fig. 6.** Subtracting residual disparity from all 2D localisations reduces 3D fit error. Insets show the magnitude of the residual disparity for each dataset. Lateral fit error is plotted on the left side of each graph and axial fit error on the right hand side as indicated. Colour bars valid for all plots.

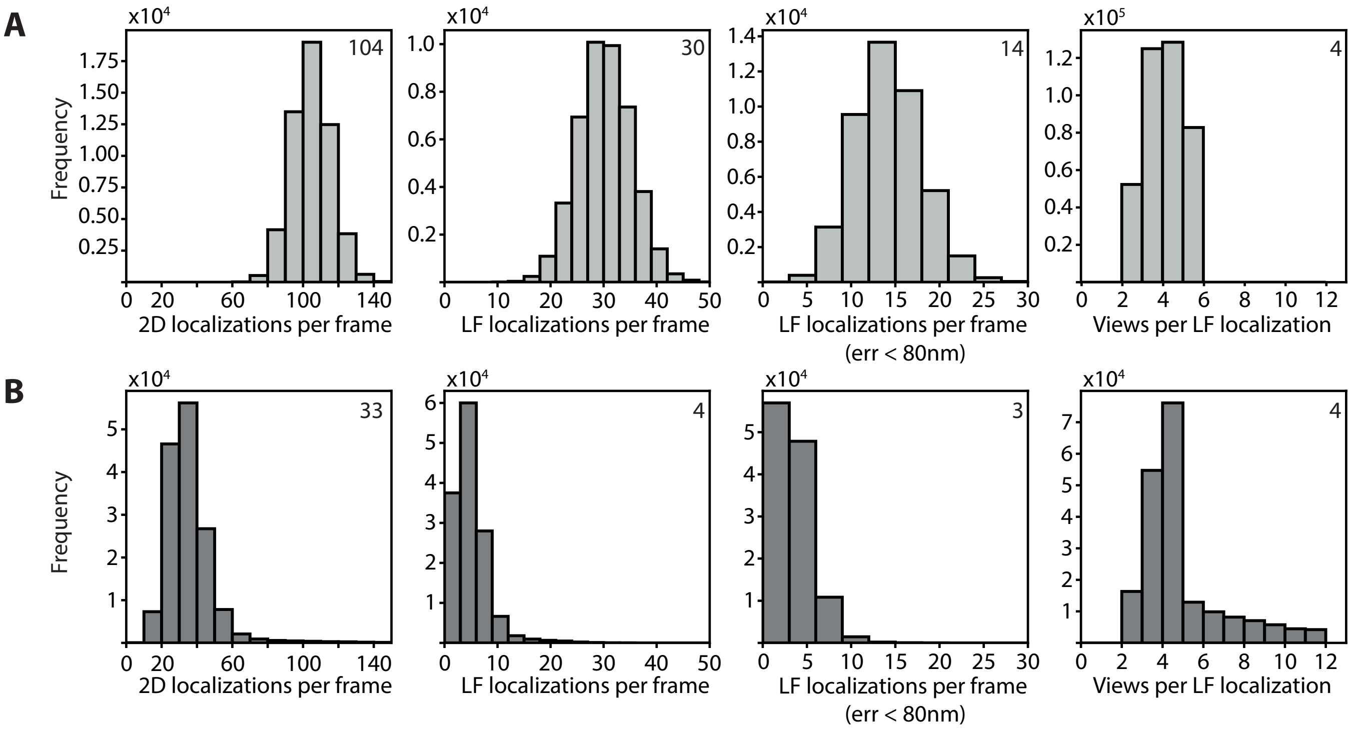


**Fig. 7.** Summary of filtering of localisations into final datasets as rendered in Figure 5 of the main text. Top row (configuration 1, (A) of Fig 5), bottom row (configuration 2, (A) of Fig 5). 2D localisations per frame refers to all localisations in all views returned by the ThunderSTORM software (15). The light field fitting algorithm returns 3D position estimates referred to as LF localisations per frame. These 3D localisations are filtered using 3D fit error as a proxy for precision to yield so-called LF localisations per frame (err < 80 nm). The final histogram in each row shows the number of views (i.e. number of 2D localisations) which contributed to each light field fit.

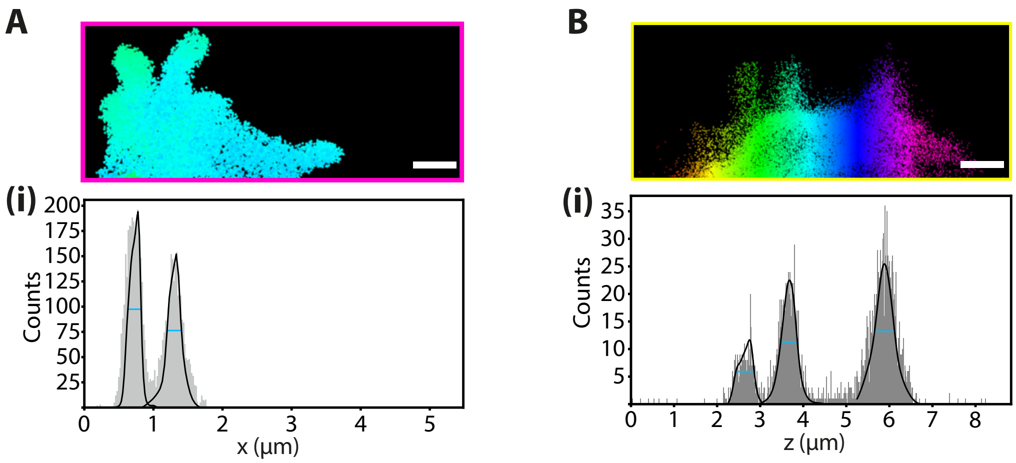


**Fig. 8.** (A) 1D histogram of data plotted in the inset of (A) (i) of Figure 5 in the main text. (B) 1D histogram of data plotted in the inset of (B) (i) of Figure 5 in the main text. Scale bars represent 1 $\mu$m.

1. R. McGorty, J. Schnitzbauer, W. Zhang, and B. Huang, "Correction of depth-dependent aberrations in 3D single-molecule localisation and super-resolution microscopy," Optics letters **39.2**, 275–278 (2014).
2. P.N. Petrov, Y. Shechtman, and W.E. Moerner "Measurement-based estimation of global pupil functions in 3D localisation microscopy," Optics express **25.7**, 7945-7959 (2017).
3. M.J. Booth, M.A. Neil, and T. Wilson "Aberration correction for confocal imaging in refractive index mismatched media," Journal of microscopy **192.2**, 90-98 (1998).
4. S.F. Gibson, F Lanni "Experimental test of an analytical model of aberration in an oil-immersion objective lens used in three-dimensional light microscopy," JOSA A **9.1**, 154-166. (1992).
5. B.M. Hanser, M. Gustafsson, M.G. Agard, J.W. Sedat "Phase-retrieved pupil functions in wide-field fluorescence microscopy" Journal of microscopy **216.1**, 32-48. (2004).
6. C.J.R. Sheppard, P. Torok, M.G. Agard, J.W. Sedat "Effects of specimen refractive index on confocal imaging" Journal of microscopy **185.3**, 366-374 (1997).
7. E.J. Botcherby, R. Juskaitis, M.J. Booth, T. Wilson "An optical technique for remote focusing in microscopy." Optics Communications **281(4)**, 880-887 (2008).
8. D. Axelrod "Evanescent excitation and emission in fluorescence microscopy." Biophysical journal **104(7)**, 1401-1409 (2013).
9. T. Ruckstuhl, J. Enderlein, S. Jung, S. Seeger "Forbidden light detection from single molecules." Optics Communications **72(9)**, 2117-2123 (2000).
10. N. Bourg, C. Mayet, G. Dupuis, T. Barroca, P. Bon, S.Lecart, E. Fort, S. Leveque-Fort "Direct optical nanoscopy with axially localized detection." Nature Photonics **9(9)**, 587 (2013).
11. M. Born and E. Wolf , "Principles of Optics." Pergamon, New York (1980).
12. M. Tokunaga, N. Imamoto and K. Sakata-Sogawa , "Highly inclined thin illumination enables clear single-molecule imaging in cells." Nature Methods **5**, 159-161 (2008).
13. C. Liesche, K. S. Grußmayer, M. Ludwig, S. Wörz, K. Rohr, D.-P. Herten, J. Beaudouin, and R. Eils, "Automated analysis of single-molecule photobleaching data by statistical modeling of spot populations," Biophysical journal **109**, 2352–2362 (2015)
14. S. Abdul Rehman, A. Carr, M. Lenz, S. Lee, and K. O'Holleran, "Maximizing the field of view and accuracy in 3D Single Molecule localisation Microscopy." Optics Express **26**, 4631-4637 (2018).
15. M. Ovesny, P. Krizek, J. Borkovec, Z. Svindrych and G.M. Hagen "ThunderSTORM: a comprehensive ImageJ plug-in for PALM and STORM data analysis and super-resolution imaging." Bioinformatics, **30**(16)2389-2390 (2014)
